# Spheroid culture of mesenchymal stromal cells results in morpho-rheological properties appropriate for improved microcirculation

**DOI:** 10.1101/440966

**Authors:** Stefanie Tietze, Martin Krater, Angela Jacobi, Anna Taubenberger, Maik Herbig, Rebekka Wehner, Marc Schmitz, Oliver Otto, Catrin List, Berna Kaya, Manja Wobus, Martin Bornhauser, Jochen Guck

**Affiliations:** Biotechnology Center, Center for Molecular and Cellular Bioengineering, TU Dresden, Tatzberg 47-49, 01307 Dresden, Germany; Medical Clinic I, University Hospital Carl Gustav Carus, TU Dresden, Fetscherstraße 74, 01307 Dresden, Germany; Institute of Immunology, Medical Faculty Carl Gustav Carus, TU Dresden, Fetscherstraßte 74, 01307 Dresden, Germany; Center for Regenerative Therapies Dresden, Center for Molecular and Cellular Bioengineering, TU Dresden, Fetscherstraßte 105, 01307 Dresden, Germany

**Keywords:** mesenchymal stromal cells, mesenspheres, cell mechanics, microcirculation mimetics

## Abstract

Human bone marrow mesenchymal stromal cells (MSCs) have been used in clinical trials for the treatment of systemic inflammatory diseases due to their regenerative and immunomodulatory properties. However, intravenous administration of MSCs is hampered by cell trapping within the pulmonary capillary networks. Here, we hypothesize that traditional twodimensional (2D) plastic-adherent cell expansion fails to result in appropriate morphorheological properties required for cell-circulation. To address this issue, we adapted a novel method to culture MSCs in non-adherent three-dimensional (3D) spheroids (mesenspheres). The biological properties of mesensphere-cultured MSCs remained identical to conventional 2D cultures. Morpho-rheological analyses revealed a smaller size and lower cell stiffness of mesensphere-derived MSCs compared to plastic-adherent MSCs, measured using real-time deformability cytometry (RT-DC) and atomic force microscopy, resulting in an increased ability to pass through micro-constrictions in an ex vivo microcirculation assay. This ability was confirmed in vivo by analysis of cell accumulation in various organ capillary networks after intravenous injection of mesensphere-derived MSCs in mouse. Our findings generally identify cellular morpho-rheological properties as attractive targets to improve microcirculation and specifically suggest mesensphere cultures as a promising approach for optimized MSC-based therapies.

---

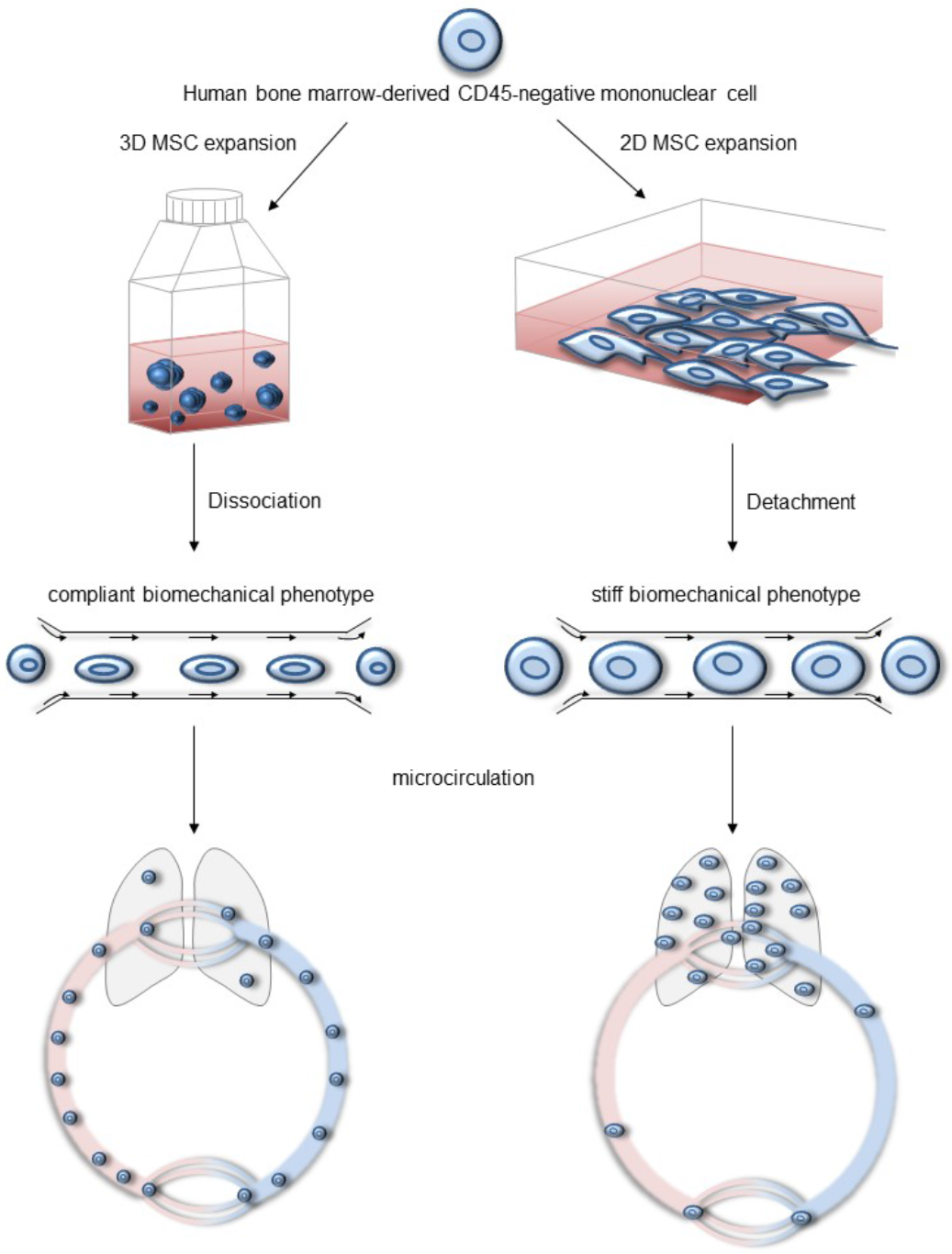

## Introduction

Mesenchymal stromal cells (MSCs) are described as a population of multipotent cells within almost all tissues, that distinguish themselves from other tissue residing cells by their unique morphology, fibroblast colony forming unit (CFU-F) capacity, multilineage differentiation potential (1, 2), and immunomodulatory properties (3). They suppress alloantigen-induced T cell proliferation (4), modulate terminal B cell differentiation (5) and inhibit the functions of natural killer cells (6) and dendritic cells (7) via the secretion of soluble factors (8, 9), and direct cell-cell interactions (10). Owing to their immunosuppressive effects and tissue regenerative potential, MSCs have been used in clinical trials testing therapies for various diseases including graft-versus-host disease after allogenic stem cell transplantation (11, 12), autoimmune diseases (13), liver diseases (14), orthopedic injuries (15), cardiovascular diseases (16), and cancer (17). For these applications, high numbers of MSCs are required, for which they are commonly isolated from tissues of healthy human donors including bone marrow (BM) explants and expanded on rigid plastic surfaces as adherent monolayer (2D) cultures. This process is associated with depletion of less adherent earlier progenitors (18). Moreover, recent work has shown that long-term culturing on rigid substrates fails to resemble the natural three-dimensional (3D) bone marrow microenvironment with its unique tissue architecture and mechanical properties. This leads to reduced growth rates, loss of multipotency, and cellular senescence (19, 20). The most eminent disadvantages, however, are alterations in the cytoskeletal organization and the morpho-rheological phenotype, such as cell size and compliance (21). Intravenous infusion of such expanded MSCs is followed by cell trapping within the pulmonary microcirculation and inefficient homing to target organs in mice (22, 23) and humans (24).

The cell’s morpho-rheology has moved into the focus of attention since recent work identified them as key regulators of cell migration (25), immune response (26) and cell polarization (27). Changes in cell deformability, especially cell softening, not only accompany cell differentiation (28) and motility they also improve cell passage through microcapillaries (29). This raises the question whether engineering MSCs with circulation-appropriate morpho-rheological properties can improve microcirculation. As a cell’s morphology and mechanical state depend highly on environmental physical properties and the resulting extracellular to intracellular signaling (30), we asked whether the modulation of morpho-rheological features can be achieved using innovative cell culture techniques. Expansion of MSCs in 3D spheroidal aggregates compared to traditional 2D techniques results in cells with smaller size (31, 32) and reduced Young’s modulus (33). Therefore, we adapted a scaffold-free 3D cell culture system for the monoclonal expansion of MSCs in non-adherent spheroids (mesenspheres). We identified that 3D-expanded cells fulfill properties of MSCs including self-renewability, multilineage differentiation and immune modulation. Based on our findings of cytoskeletal rearrangement in 3D-expanded MSCs, we compared morpho-rheological properties of MSCs after scaffold-free 3D and traditional 2D expansion. Using atomic force microscopy (AFM) indentation measurement and real-time deformability cytometry (RT-DC), we found that 3D-expanded MSCs are smaller and more compliant. Ex vivo microcirculation analysis revealed that mesensphere-forming single cells compared to plastic-adherent MSCs required less time to enter and pass a microfluidic microcirculation mimetic (MMM) device. Furthermore, we demonstrated that after intravenous injection of MSCs in the tail vein of mice, mesensphere-derived MSCs were detected to a larger amount in the microcirculation of various organs other than the lungs. In this respect, the unique morpho-rheological properties of mesensphere-expanded MSCs make them amenable for the development of advantageous stem cell therapies to overcome limitations that arise with traditional 2D cultures.

## Results

### Mesenspheres comprise multipotent, self-renewable and immunomodulatory MSCs capable of hematopoietic stem and progenitor cell support

Mesenchymal stromal cell spheroids (mesenspheres) were cultured from bone marrow (BM) mononuclear cell (MNC) fraction isolated by density gradient centrifugation. Enrichment of MSCs was achieved by immunomagnetic depletion of CD45-positive MNCs. Utilizing ultra-low attachment flasks and mesensphere growth medium, CD45-depleted MNCs formed self-assembled spherical-shaped mesenspheres. Hematoxylin and eosin (H&E) staining of paraffin-embedded sections revealed that mesenspheres build a compact 3D network composed of small round cells surrounded by elongated flat cells (Figure 1A). To determine the frequency of mesenchymal progenitors, we performed CFU-F assay. Mesensphere-derived MSCs were found to establish multiple fibroblast colonies with various sizes. Using limiting dilution analysis (ELDA) (34) the estimate of stem cell frequency in mesenspheres was 1/82.7 cells with 95% confidence interval of 1/50.9 to 1/134 cells (Figure 1B). According to the minimal criteria for MSCs (35), mesensphere cells were found positive for surface marker expression of CD105 (85.5% ± 8.5%), CD73 (94.3% ± 5.7%), and CD90 (85% ± 16%) and lack expression of CD14 (4.6% ± 1.2%), CD19 (2.2% ± 2%), CD34 (2.5% ± 1.6%), CD45 (4.2% ± 5%) and HLA-DR (13.4% ± 12.4%; Figure 1C). Moreover, we found that mesenspheres exhibit multilineage differentiation potential (Figure S1A-E). Immunomodulatory properties of mesensphere-derived MSCs were investigated using a modified mixed lymphocyte reaction assay (36). Lymphocyte proliferation as measured by [^3^H]-thymidine incorporation was significantly decreased in the presence of mesensphere MSCs (44097.7 cpm ± 23200.0 vs. 63718.3 cpm ± 32461.1; p = 0.0044; Figure 1D). Furthermore, BM-derived MSCs have been shown to support hematopoietic stem and progenitor cell (HSPC) maintenance and engraftment (37). We found that clonogenic HSPCs with appropriate differentiation potential could be efficiently expanded in co-culture with mesenspheres (Figure S2A-G).

**Figure 1:**
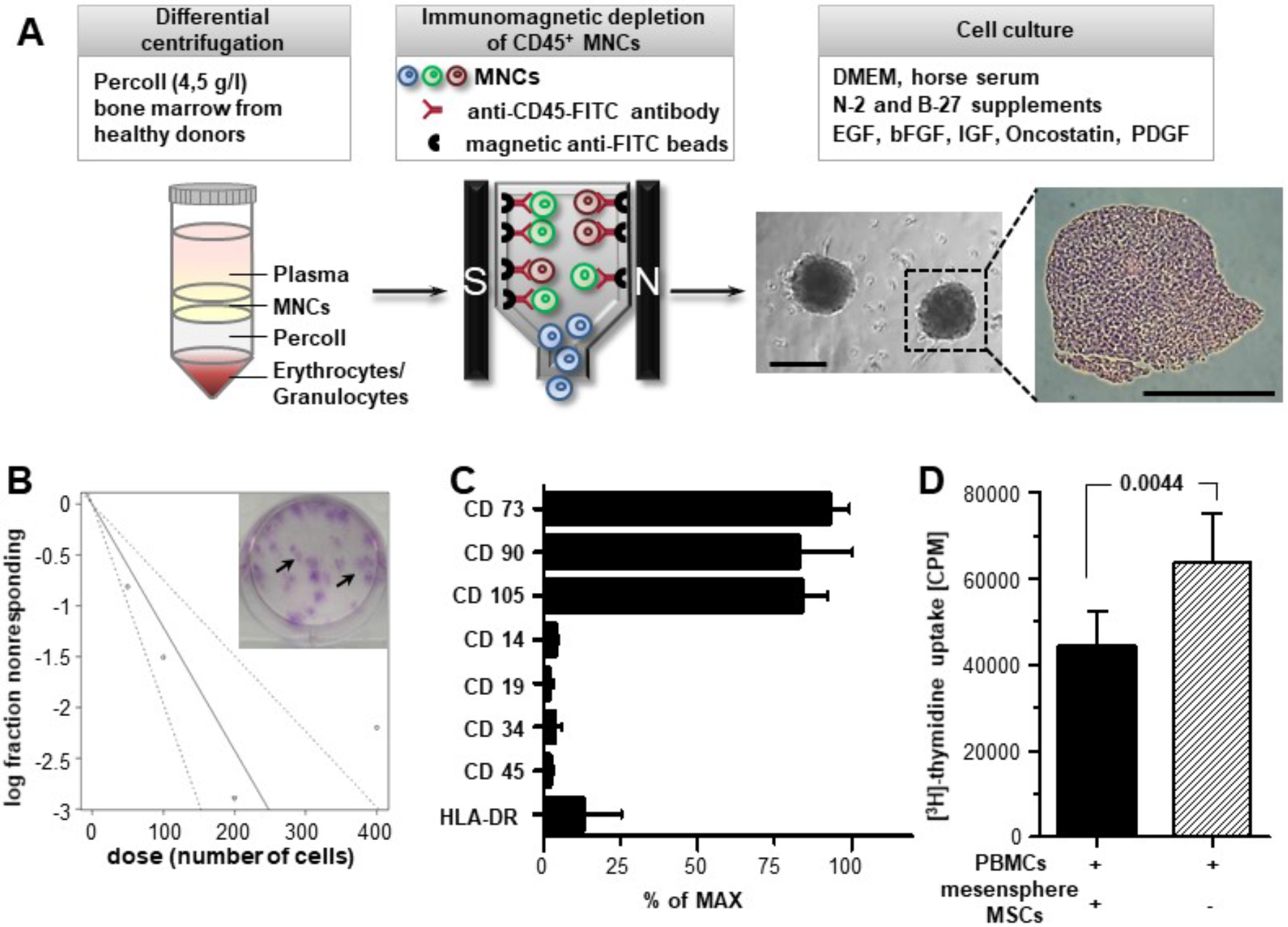
Mesenspheres comprise multipotent, self-renewable and immunomodulatory MSCs. (A) Cell culture of mesenspheres, MSCs were isolated from bone marrow of healthy donors by density centrifugation and immunomagnetic depletion of CD45-positive mononuclear cells (MNCs). Representative hematoxylin and eosin (H&E) staining of paraffin-embedded mesensphere sections. Scale bar, 200 μm. (B) Extreme limiting dilution assay (ELDA) of mesensphere MSCs, after a 2-week culture period mesensphere MSCs were plated at a density of 5,10, 20 and 40 cells cm^2^. The frequency of mesenchymal progenitors was calculated using ELDA method by counting fibroblast colonies in each dilution of three independent experiments including three technical replicates. Representative picture showing colony-forming unit fibroblast (CFU-F) assay using mesensphere MSCs at a clonal density of 40 cells cm^2^. (C) Flow cytometry analyses of mesensphere MSCs for minimal criteria cell surface marker expression (CD73+, CD90+, CD105+, CD14−, CD19−, CD34−, CD45−, HLA-DR). Histogram bars representing mean ± s.e.m. of four independent replicates. (D) Modified mixed lymphocyte reaction assay, histogram representing [^3^H]-thymidine incorporation into CD3/CD28-stimulated peripheral blood mononuclear cells (PBMCs) after co-culture with irradiated (30 Gy) mesensphere MSCs (1/30 (PBMC/MSC) ratio). Histogram bars representing mean ± s.d. of five independent experiments. Statistical significance was determined by an unpaired two-tailed t-test.

Together, these results show that the biological properties of mesensphere-derived MSCs are identical to those cultured conventionally on rigid 2D plastic surfaces. For their clinical use, MSC expansion generally relies on 2D culturing techniques to reach relevant numbers. However, after intravenous administration of such expanded MSCs cell delivery to target tissues is limited as a result of mechanical cell trapping within the pulmonary capillary networks

### Mesensphere-derived MSCs exhibit characteristic morpho-rheological properties

It has been shown that the size and cytoskeletal proteins of MSCs maintained in 3D (mesenspheres) are drastically reduced compared to 2D (31, 38). Therefore, we studied whether mesensphere culture of MSCs also affects other relevant morpho-rheological properties that can influence microcirculation. Analysis of the cytoskeleton distribution in entire mesenspheres revealed short, tightly bundled actin filaments in the apical mesensphere region. Towards the mesensphere interior actin filaments assembled as cell cortex associated ring-like belt structures (Figure 2A). In contrary, in plastic-adherent MSCs, we observed uniformly distributed actin stress fibers, namely dorsal stress fibers, transverse arcs and ventral stress fibers (Figure 2B). Since cytoskeletal mutations directly correlate with morphorheological properties (29, 39, 40), we performed atomic force microscopy (AFM) indentation measurements on single MSCs in entire mesenspheres and MSCs cultured on rigid plastic surfaces to probe cell stiffness (Figure 2C). The morpho-rheological properties were quantified by extracting the apparent Young’s modulus from the obtained force-indentation curves. As demonstrated in Figure 2D, apparent Young’s moduli measured for mesenspheres were significantly lower compared to plastic-adherent MSCs (761.4 Pa ± 542.4 vs. 5926 Pa ± 486.9, p = 4.9×10^−8^). Interestingly, the apparent Young’s moduli of mesensphere cells that had been migrated out of mesenspheres after 5 hours were increased compared to cells within mesenspheres (1871 Pa ± 971.1 vs. 761.4 ± 542.4, p = 0.001), which identifies plastic adherence as a main contributor of cell mechanics. Furthermore, morpho-rheological properties of suspended MSCs were analyzed using real-time deformability cytometry (RT-DC) (Figure 2E). Here, the apparent Young’s modulus can be derived from image analysis that quantified cell size and the resulting deformation as cells pass through a narrow microfluidic channel, similar to blood flow, in real-time and with high-throughput (up to 1,000 cells/sec) (39, 41, 42). In comparison with plastic-adherent MSCs, mesensphere-derived MSCs appeared significantly smaller (cross-sectional area: 290,5 μm^2^ ± 34,9 vs. 391,02 μm^2^ ± 30,4, p = 0.0008, Figure 2F) and more deformable (deformation: 0.056 ± 0.006 vs. 0.041 ± 0.004, p = 0.0166, apparent Young’s modulus: 1129.6 ± 135.0 Pa vs. 1812.6 Pa ± 98.0, p = 0.0002, Figure 2G-H). So far, it has been assumed that the large size of MSCs cultured on rigid 2D plastic surfaces causes trapping within the pulmonary capillaries (23). However, our previous studies indicate that in addition to cell size, also the cell mechanical properties affect microcirculation (29). Therefore, we hypothesized that altered morpho-rheological properties of MSCs cultivated in mesenspheres can improve microcirculation.

**Figure 2:**
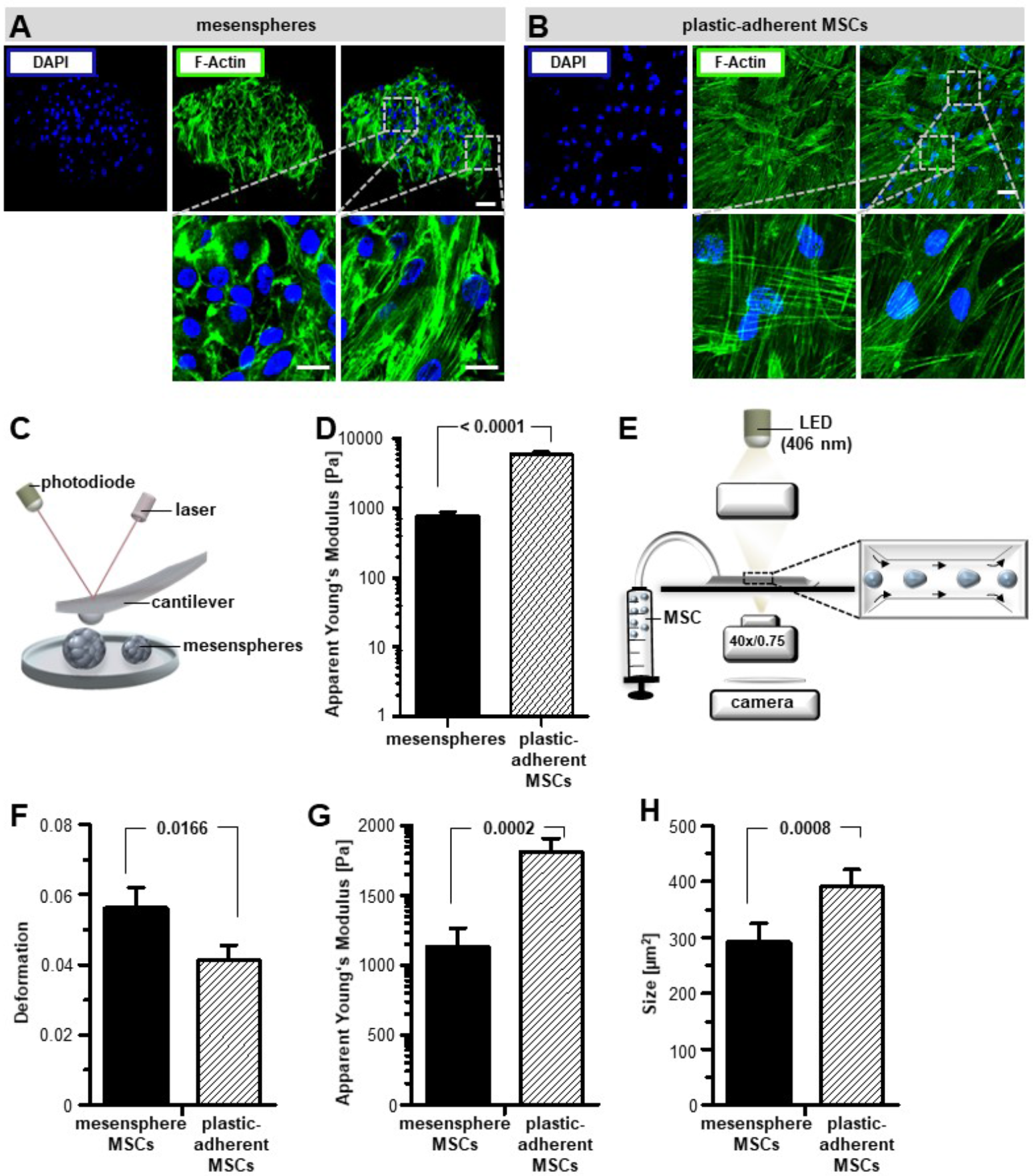
Mesensphere-derived MSCs exhibit exceptional morpho-rheological properties. (A-B) Representative confocal microscopy maximal projections (Z = 15 μm) of (A) mesensphere cytoskeletal structures and (B) plastic-adherent MSC cytoskeletal structures via staining of filamentous actin (F-Actin, green) and nuclei (blue). Scale bars upper panels, 65 μm; Scale bars lower panels, 15 μm (C) Schematic presentation of atomic force microscopy (AFM) measurement of cell stiffness by indentation at mesenspheres and plastic-adherent MSCs. The mechanical properties of the cells were quantified using the apparent Young’s modulus calculated from the obtained force-indentation curves. (D) Histogram bars represent mean ± s.e.m. of two independent atomic force microscopy measurements including nine technical repeats. Statistical significance was determined using an unpaired two-tailed t-test. (E) In real-time deformability cytometry (RT-DC) mesensphere MSCs and plastic-adherent MSCs were flowed through a 30 μm × 30 μm microfluidic constriction and were deformed without contact by shear stress and pressure gradients. (F-H) Cellular morpho-rheological properties of mesensphere MSCs and plastic-adherent MSCs were quantified as (F) cell deformation, (G) apparent Young’s Modulus, and (H) cell size (cross-sectional area) using automated image analysis. Histogram bars represent mean ± s.d. of four independent experiments. Statistical significance was determined using a 1-dimensional linear mixed model and a likelihood ratio test.

### Morpho-rheological properties of mesensphere-derived MSCs permit cell circulation

To analyze the impact of the culture system, and the resulting altered morpho-rheological properties, on MSC circulation, we mimicked pulmonary microcirculation in a microfluidic device, namely microfluidic microcirculation mimetic (MMM). MMM consists of a serpentine microchannel 300 μm in total length and 20 μm in height and 20 μm in width with 187 successive constrictions smaller than the cell diameter (5 μm in cross section; Figure 3A) (43, 44). Thus, the constrictions and therefore the cell deformations occur sequentially one after the other as in the microcirculation. Cells were re-suspended in PBS and flushed through the MMM using a constant pressure difference of 150 mBar between the inlet and the outlet. To quantify the ability of cells to pass through the constrictions, we measured entry time (time that is needed to pass the first constriction of the microchannel) and total passage time. Mesensphere MSCs were found to require less time to enter the first constriction (0.035 sec ± 0.0127 vs 0.106 sec ± 0.0261, p = 0.0077) and less time to pass the whole microchannel (0.470 sec ± 0.0167 vs 1.384 sec ± 0.0778, p < 0.0001) compared to plastic-adherent MSCs (Figure 3B, C). We therefore concluded that, in addition to cell morphology, also physical deformability affects microcirculation of MSCs. Interestingly, we found that 2D expanded MSCs rapidly change their morpho-rheological phenotype when cells were detached and grown in non-adherent 96-well plates. Here, cells stick together and form spheroidal aggregates (secondary spheres) within one day (15,000 cells per sphere). After dissociation of secondary spheres, we obtained MSCs with morpho-rheological properties (deformation and size) similar to those expanded in mesenspheres (Figure S3A-C). Moreover, these cells compared to plastic-adherent MSCs were also characterized by improved circulation through MMM with 8 μm in cross section (Figure S3D). These findings confirm that morpho-rheological phenotype of MSCs can be manipulated by the appropriate cell culture technique.

**Figure 3:**
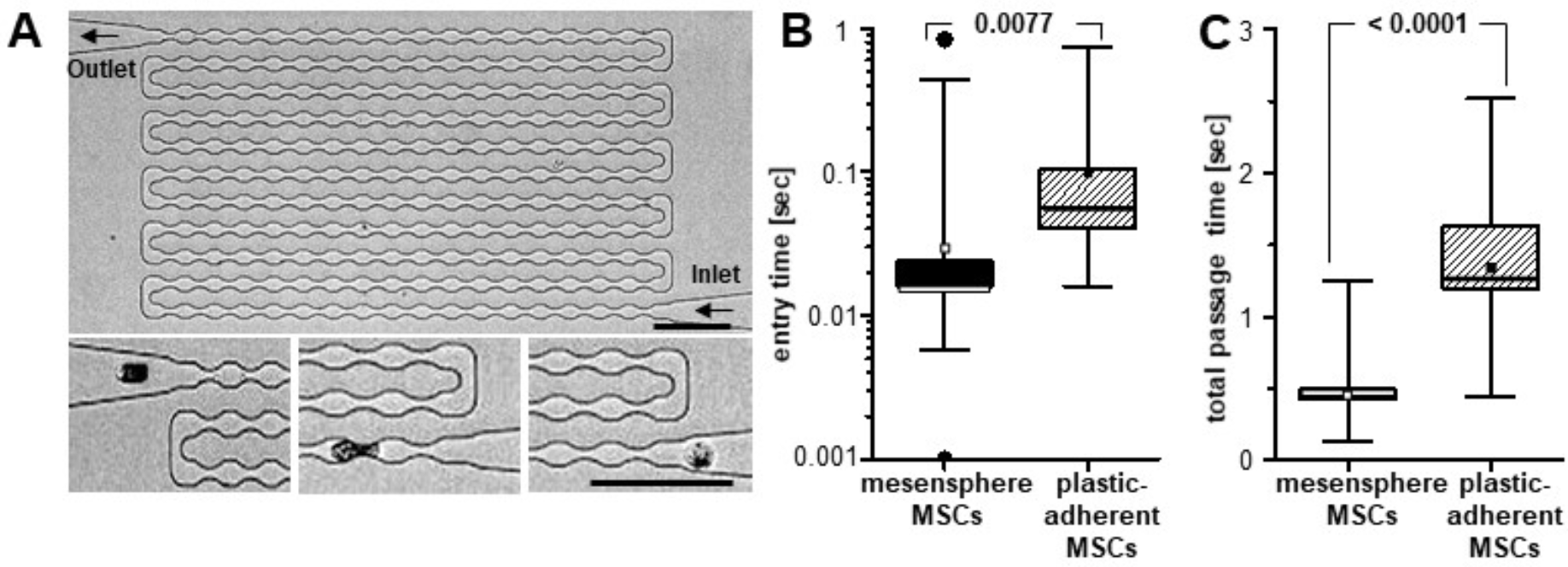
Ex vivo microfluidic microcirculation mimetic analysis of mesensphere-derived and plastic-adherent MSCs. (A) Phase contrast image of a MMM with 187 successive constrictions (with 5μm smallest in width). Smaller inlets show the close-up of a cell before the first constriction, after the first two constrictions, and after the last constriction (from right to left). Scale bars are 100 μm. (B) Box & whiskers blots representing entry time (passage through the first constriction) and (C) total passage time (from inlet to outlet) for mesensphere MSCs and plastic-adherent MSCs to enter and pass a MMM with successive constrictions. Lines representing mean ± s.d. of two independent experiments including thirty technical repeats. Statistical analyses were determined using an unpaired two-tailed t-test.

To further investigate whether the unique morpho-rheological properties of mesensphere-derived MSCs promote increased microcirculation, MSCs derived from mesenspheres or plastic-adherent cultures were dissociated to obtain single-cell suspensions and injected intravenously into the tail vein of NSG mice (Figure 4A). After 15 min mice were sacrificed, organs were dissected, and DNA was isolated to quantify human Alu-sequences within tissues. As depicted in Figure 4B, real-time PCR for human Alu-sequences revealed that mesensphere MSCs can pass lung capillaries more efficiently resulting in a more even distribution within the capillary networks of other tissues. Compared to 2D cultured MSCs, lung trapping of mesensphere MSCs decreased by about 30%, whereas recovery in liver, heart, spleen, and kidney increased up to 20% (Figure 4C-F). These findings demonstrate that efficient microcirculation of MSCs is determined by their morpho-rheological phenotype. This microcirculation appropriate morpho-rheological phenotype can be engineered using 3D mesensphere instead of 2D plastic adherent cultur techniques and thus might be crucial for effective future MSC-based therapies.

**Figure 4:**
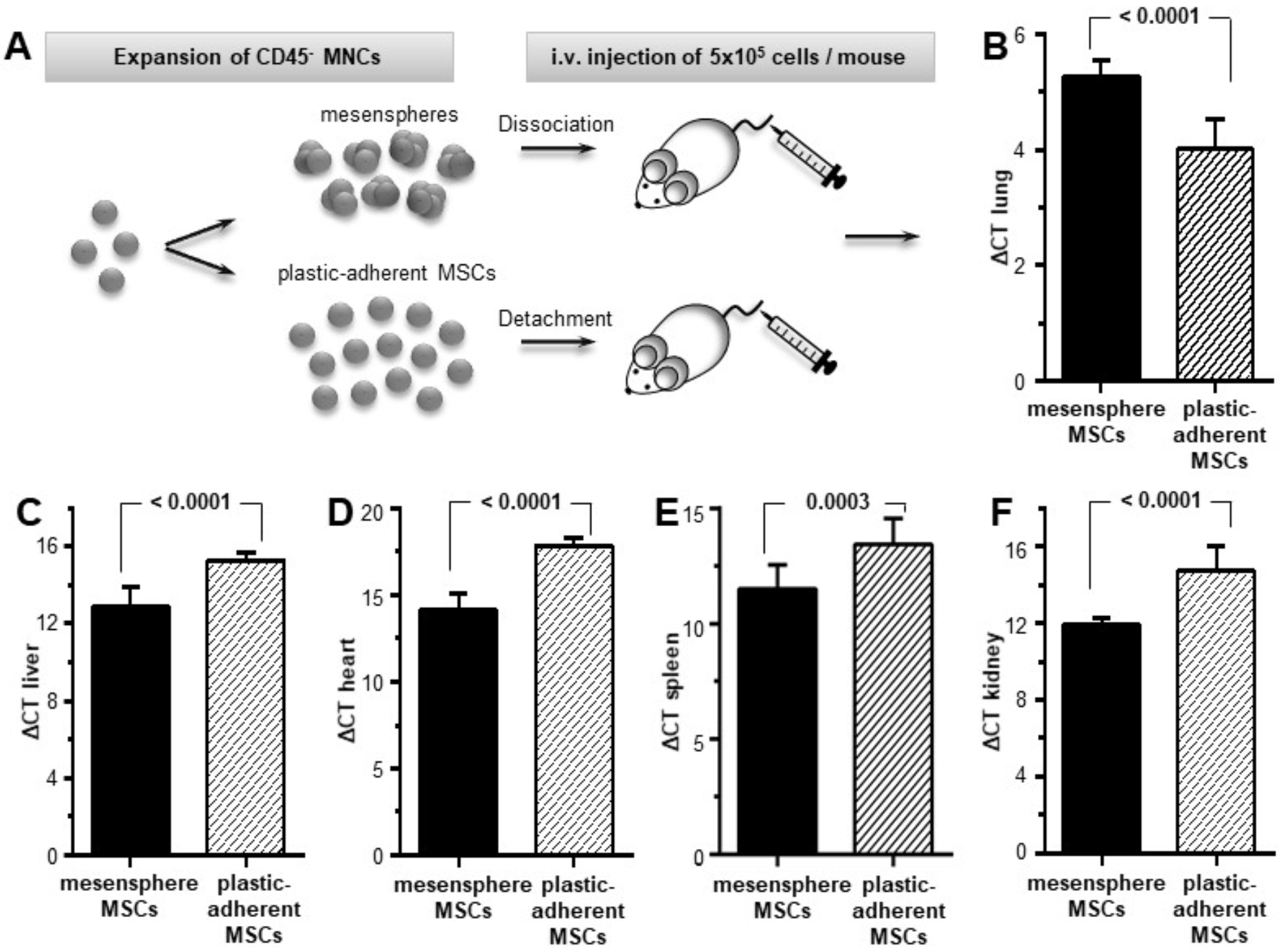
Microcirculatory properties of mesensphere and plastic-adherent expanded MSCs. (A) For analyses of microcirculatory properties 5×10^5^ MSCs from mesenspheres or 2D cultures were injected intravenously into the tail vein of NSG mice. Human MSCs were detected via amplification of human Alu sequences 15 min after i.v. infusion. (B-F) Histogram bars representing amplification of human Alu sequences normalized to glyceraldehyde-3-phosphate-dehydrogenase GAPDH (ΔCT) in (B) lung, (C) liver, (D) heart, (E) spleen, and (F) kidney 15 min after i.v. infusion of mesensphere MSCs in comparison to plastic-adherent MSCs. Histogram bars representing mean ± s.d. of twelve independent experiments. Statistical analyses were determined using an unpaired two-tailed t-test.

## Discussion

Since Friedenstein et al. first described MSCs as a clonogenic population of BM-derived cells (45), they have been applied in numerous clinical trials due to their regenerative and immunomodulatory potential. However, prior to the use of MSCs as cell-based therapeutics, considerable hurdles must be overcome such as establishment of standards for cell purification and expansion (46). These standards include conventional 2D monolayer cultures on rigid tissue culture plastic, which has been linked to loss of MSC function (19–21). Therefore, it is crucial to develop physiologically relevant culture environments that maintain MSCs in a more naïve state.

Méndez-Ferrer et al. adapted a culture regime, conducive for the assembly of Nestin+ cells into neurospheroids, for Nestin+ murine and human BM MSCs to obtain mesenspheres (47). Using slightly modified culture conditions, we succeeded to obtain mesenspheres derived from CD45-depleted BM-MNC. It is supposed that spheroid generation on low attachment plates starts with loose cell aggregates mediated by ECM-integrin interactions followed by cell compaction via cadherin bindings (38, 48). The scaffold-free culture of mesenspheres yielded a highly heterogeneous cell population in cell size and cell morphology. Irrespectively, mesensphere forming cells behave functionally like MSCs as shown by their CFU-F capacity, cell surface molecule expression, multilineage differentiation, and HSPC support. Previous studies reported that cell expansion in 3D spheroids even enhances the differentiation potential of multipotent cells (49). Moreover, it is presumed that the 3D cell culture environment improves the therapeutic potential of MSCs by increased expression of relevant genes such as CXCR4 promoting adhesion to endothelial cells (50) or SCF that is involved in HSPC maintenance (51). Our analysis demonstrated that mesenspheres express high levels of HSPC-supporting factors and are capable to expand a population containing CD45+ / CD34+ / CD133+ long-term HSPC. Hence, efficient expansion of functional MSCs does not require plastic adherence. MSCs expansion can be achieved using scaffold-free 3D cell culture techniques that mimic the physiological BM microenvironment more closely than plastic-adherent 2D monolayer cultures.

Since MSCs can modulate multiple components of the immune system including T- and B-cell proliferation, they emerge as promising candidates for cell-based immunotherapies. Our analysis revealed that mesensphere forming cells effectively suppress T cell proliferation in the same manner as shown for clinical used MSCs grown in 2D monolayers (4). This suggests that mesensphere MSCs possess immunomodulatory properties that qualifies them as therapeutics for systemic inflammatory diseases such as graft-versus-host disease (GvHD) after allogenic hematopoietic stem cell transplantation.

So far, the most convenient route of administration for traditional 2D-expanded MSCs is intravenous injection. Undesirably, homing to target organs is often impeded by cells trapped within the pulmonary microcapillary networks (22–24). Previous studies with neutrophils revealed that stiff primed compared to soft resting neutrophils exhibit longer total passage times in microcirculatory mimetics (44). Moreover, recent clinical observations provide evidence that primed neutrophils are retained within the pulmonary microcirculation, while deprimed neutrophils are released into the systemic circulation (52). Lautenschläger et al. found that mobility of neutrophils is facilitated by cell softening, which appeared to be regulated by the actin cytoskeleton (25, 29). This implies that cell deformability must be crucial for efficient passage kinetics in microcirculation. Notably, mesensphere compared to 2D cultured MSCs appeared smaller and more deformable. These results agree with previous studies in which plastic-adherent MSCs are described as a little-deformable population after expansion (53). The altered morpho-rheological properties of mesensphere MSCs are even prominent in suspension after enzymatic dissociation of mesenspheres. These results may suggest that mesensphere MSCs exhibit improved microcirculatory properties. Our assumption was confirmed by ex vivo microcirculation analyses, in which mesensphere forming cells compared to plastic-adherent MSCs required less time to enter and pass a microfluidic device. This is in line with findings by Chan et al. that showed a clear correlation between cell stiffening and extended total passage time in human promyelocytic leukemia cells (HL60). Moreover, they found that inhibition of actin polymerization with cytochalasin D decreases total passage time (43), which indicates that increased actin polymerization in plastic-adherent MSCs compared to mesensphere cultured cells modulates cell stiffness and therefore might increase total passage time. The results obtained by MMM analyses were corroborated by *in vivo* experiments, where mesensphere MSC derived DNA was detected to a larger amount in organ microcirculation other than the lung capillaries after intravenous injection into the tail vein of NSG mice. The differences became particularly evident in the liver, heart, spleen, and kidney. These findings presumably predict an increased homing of MSCs to target organs. In combination with the reported immunomodulatory properties of mesensphere MSCs these findings will be of considerable clinical significance for the treatment of systemic inflammatory diseases.

Thus, in future it might be promising to use non-adherent 3D reconstitution techniques for the development of MSC cell-based therapeutic approaches in order to ensure expansion of MSCs with small size and enhanced deformability compared to plastic-adherent MSCs. This will prevent them from getting retained in the lung — the first major organ with an extensive microcirculatory network that MSCs have to pass — and contribute to a more even distribution of MSCs in organ capillary networks after intravenous injection. Therefore, using innovative cell culture techniques modulating the morphological and rheological phenotype of MSCs will significantly increase the feasibility of clinical stem cell therapeutic approaches.

## Material and methods

### Cell isolation and culture

MSCs were isolated from healthy volunteer donors after obtaining informed written consent (ethical approval no. EK221102004, EK47022007) according to modifications of a previously reported method (54, 55). Briefly, human bone marrow aspirates were diluted in PBS (Thermo Fisher Scientific, Waltham, USA) at a ratio of 1:5. A 20 ml aliquot was layered over a 1,073 g/ml Percoll solution (Biochrom, Berlin, Germany) and centrifuged at 2000 × g for 15 min at room temperature. Mononuclear cell (MNC) fraction was recovered, washed twice in PBS and stained with anti-CD45-FITC antibodies. For immunomagnetic enrichment anti-FITC magnetic beads (Miltenyi Biotec GmbH, Bergisch Gladbach, Germany) were used according to the manufacturer’s instructions.

For clonal sphere formation 5 × 10^6^ cells were seeded in ultra-low attachment culture flasks (Greiner, Kremsmuenster, Austria). The mesensphere growth medium contained 15% horse serum, 2% B27 and 1% N2 supplements, epidermal growth factor (EGF), fibroblast growth factor-basic (bFGF), insulin-like growth factor (IGF; 20 ng ml^−1^), oncostatin and platelet-derived growth factor (PDGF; 10 ng ml^−1^) in DMEM/F12, HEPES/human endothelial-SFM (Thermo Fisher Scientific, Waltham, USA). The cultures were kept at 37 °C with 5% CO_2_ in a water-jacked incubator. For secondary sphere formation plastic adherent MSCs were detached and 15,0 cells were seeded per well of a 96-well plate pre-coated with 15 μl 1% agarose in DMEM + 10% FCS. Medium changes were performed weekly. For further experiments mesenspheres were enzymatically dissociated in DMEM, high glucose supplemented with 2.2% GlutaMAX (Thermo Fisher Scientific, Waltham, USA), 0.2% type I collagenase (Wako Chemicals GmbH, Neuss, Germany), and 2.5% type I DNase (Sigma-Aldrich, St. Louis, USA). Mobilized peripheral blood was purchased from healthy donors treated with granulocyte colony-stimulating factor (G-CSF; 7.5 μg kg^−1^) for 5 days after obtaining informed written consent (ethical approval no. EK221102004, EK47022007). CD34^+^ HSPCs were purified from leukapheresis using anti-CD34 magnetic beads (Miltenyi Biotec GmbH, Bergisch Gladbach, Germany) according to the manufacturer’s instructions. Up to 2 × 10^4^ CD34+ HSPCs were co-cultured with mesenspheres in low-cytokine HSPC growth medium on agarose-coated 24-well plates at 37 °C with 5% CO_2_ in a water-jacked incubator. Low-cytokine HSPC growth medium contained CellGro (CellGenix GmbH, Freiburg, Germany) supplemented with interleukine-3 (IL-3), stem cell factor (SCF) and FMS-like tyrosine kinase-3 (Flt-3; 2.5 ng ml^−1^; Miltenyi Biotec GmbH, Bergisch Gladbach, Germany) on agarose-coated 24-well plates at 37 °C with 5% CO_2_ in a water-jacked incubator.

### *In vitro* differentiation

Osteogenesis and adipogenesis were induced by culturing MSCs with the respective differentiation medium for 14 days, as described previously(56). Briefly, osteogenic differentiation medium contained 10 mM β-glycerophosphate, 1.5 μM dexamethasone (Sigma-Aldrich, St. Louis, USA), 0.2 mM L-ascorbate (Synopharm GmbH, Barsbuettel, Germany) and 10% FCS in DMEM (Thermo Fisher Scientific, Waltham, USA). Adipogenic differentiation was induced with 1 nM dexamethasone, 500 nM 3-isobutyl-1-methylxanthine, 100 nm indomethacin, 1 μg ml^−1^ insulin (Sigma-Aldrich, St. Louis, USA) and 10% FCS in DMEM. All cultures were kept at 37 ° C with 5% CO_2_ in a water-jacked incubator. Medium changes were performed weekly.

To quantify osteogenic differentiation potential alkaline phosphatase (ALP) activity was measured as described previously(57). Briefly, cells were lysed in 1.5 mM Tris, pH 10, containing 1 mM ZnCl_2_, 1 mM MgCl_2_ and 1% Triton X-100. Lysates were clarified by centrifugation and aliquots incubated with 3.7 mM 4-nitrophenylphosphate in 100 mM diethanolamine, pH 9.8, containing 0.1% Triton X-100 for 30 minutes at room temperature. The reaction was stopped with 100 mM NaOH and the release of 4-nitrophenolate measured photometrically at 405 nm. In alignment with the protein concentration ALP activity was calculated using a standard curve prepared with p-nitrophenol.

For histological visualization of mesenchymal lineages cells were washed with PBS and fixed with 4 % paraformaldehyde (Merck KGaA, Darmstadt, Germany) in PBS. Cells differentiated towards osteogenic lineage were detected by van Kossa staining. Briefly, fixed cultures were stained with 1% silver nitrate (Sigma-Aldrich, St. Louis, USA), developed using 0.1% pyrogallol and stabilized by 0.5% sodiumthiosulfate (Merck KGaA, Darmstadt, Germany). Adipogenic differentiation was assessed by Oil Red O staining. Concisely, fixed cells were stained with 0.1% Oil Red O solution (Sigma-Aldrich, St. Louis, USA), followed by washes with distilled water.

### Colony-forming unit-fibroblast (CFU-F) assay

Mononuclear cells were seeded in NH Expansion Medium (Miltenyi Biotec GmbH, Bergisch Gladbach, Germany) at a surface area density of 4 × 10^4^ cells cm^−2^. To determine frequency of mesenchymal progenitors in mesenspheres, serial dilutions of mesensphere-derived MSCs ranging from 50 to 400 cells in NH Expansion Medium were plated in 6-well plates. The cultures were kept at 37 °C with 5% CO_2_ in a water-jacked incubator. At day 14 cells were stained with Giemsa staining solution (Merck KGaA, Darmstadt, Germany) and adherent colonies were counted.

### Colony-forming unit (CFU)-GEMM assay

Up to 5 × 10^2^ CD34^+^ HSPCs were cultivated in 1 ml MethoCult GF + H4435 (StemCell Technologies, Vancouver, Canada) for 14 days at 37 °C in a water-jacked incubator containing 5% CO_2_. The number of colony forming units (erythroid burst-forming unit (BFU-E), colony-forming unit granulocyte (CFU-G), colony-forming unit monocyte (CFU-M), colony-forming unit granulocyte/monocyte (CFU-GM) and colony-forming unit granulocyte/erythrocyte/monocyte/megakaryocyte (CFU-GEMM)) were enumerated by optical and morphological properties using light microscopy (Axiovert 25 microscope, Carl Zeiss AG, Oberkochen, Germany).

### Flow cytometry

The following antibodies were used: anti-CD45-PECy7 (BD, Franklin Lakes, USA), anti-CD90-APC (eBioscience Inc., San Diego, USA), anti-CD105-APC, anti-CD34-PE, anti-HLA-DR-APC, anti-CD73-PE, anti-CD14-APC, anti-CD19-APC, anti-CD133-FITC (Miltenyi Biotec GmbH, Bergisch Gladbach, Germany). To evaluate proliferation and apoptosis carboxyfluorescein diacetate succinimidyl ester (CFSE) staining (Thermofisher Scientific, Waltham, USA) and propidium iodide (PI) staining (eBioscience Inc., San Diego, USA) were performed, respectively, according to the protocols of the manufacturers. Multiparameter stained cell analyses were performed using MACSQuant (Miltenyi Biotec GmbH, Bergisch Gladbach, Germany) and analyzed by FlowJo software version 7.6.5 (TreeStar Inc., Ashland, USA).

### Immunomodulatory assay

For immunomodulatory assay a modified mixed lymphocyte reaction (MLR) was performed as described previously (36). Briefly, peripheral blood mononuclear cells (PBMCs) were isolated from healthy volunteer donors after obtaining written consent (ethical approval no. EK206082008). Peripheral blood samples were diluted in PBS at a ratio of 1:2. A 20 ml aliquot was layered over a 1.077 g ml^−1^ Biocoll solution (Biochrom, Berlin, Germany) and centrifuged at 1500 × g for 30 min. MNC fraction was recovered, washed twice with PBS and resuspended in RPMI 1640 supplemented with 10% FCS (Thermo Fisher Scientific, Waltham, USA). MLR was performed by mixing 1 × 10^4^ PBMCs, CD3/CD28 Dynabeads (Life Technologies, Carlsbad, USA) and 3 × 10^3^ irradiated (30 Gy, Gammacell 3000 Elan device, Best Theratronics Ltd., Ottawa, Canada) mesensphere MSCs or plastic-adherent MSCs. After 4 h incubation at 37 °C with 5% CO_2_ in a water-jacked incubator, 1 μCi [^3^H]-thymidine (Hartmann Analytic, Braunschweig, Germany) was added to the culture. After 18 h, cells were harvested by using the Inotech Cell Harvester (Inotech Biosystems International Inc., Derwood, USA). ^3^[H]-thymidine incorporation was determined with the 1450 MicroBeta TriLux (PerkinElmer Inc., Waltham, USA), converting degree of radioactivity to counts per minute (cpm).

### Immunostaining and Histology

For confocal laser scanning microscopy, mesenspheres or plastic-adherent MSCs were fixed with 4% paraformaldehyde in PBS for 20 min. Fixed cells were stained with DAPI (Sigma Aldrich, St. Louis, USA) and phalloidin-AF488 (Molecular Probes, Eugene, USA) according to the protocols of the manufacturers. Subsequently, specimens were cover slipped in a drop of mounting medium (Polysciences Inc., Warrington, USA) and examined by a confocal laser scanning microscope (LSM 510 Meta, Leica, Wetzlar, Germany).

For histological analysis, mesenspheres were embedded in paraffin for sectioning. Microtome sections were processed with routine hematoxylin and eosin (H&E) staining and images were acquired using Axiovert 25 microscope (Carl Zeiss AG, Oberkochen, Germany).

### Atomic force microscopy (AFM)

Mesenspheres or 1 × 10^4^ plastic-adherent MSCs in CO_2_ independent medium (Life Technologies, Carlsbad, USA) were plated into round (diameter 35 mm) cell-culture dishes (FluoroDishTM, WPI, Sarasota, USA) that had been coated with 5 μg cm^−2^ fibronectin (Roche, Germany). AFM measurements were performed on a Nanowizard 4 (JPK Instruments, Berlin, Germany) equipped with a Petridishheater at 37 °C in a heat-controlled chamber. Arrow-T1 cantilevers (Nanoworld, Neuchatel, Switzerland) were modified with a polystyrene bead (radius 5 μm, microparticles GmbH, Berlin, Germany) to obtain a well-defined indenter geometry and decrease local strain during indentation. Prior to experiments cantilevers were calibrated using built-in procedures of the SPM software (JPK Instruments, Berlin, Germany). The cantilever/bead was positioned over single mesenspheres or plastic-adherent MSCs, lowered at a defined speed (5 μm sec^−1^) until a setpoint of 2.5 nN was reached and retracted. During the force-distance cycle, the force was recorded for each piezo position. The resulting force-distance curves were transformed into force- versus- tip sample separation curves and fitted with the Hertz/Sneddon model for a spherical indenter (58, 59) using the JPK analysis software (JPK DP, JPK Instruments, Berlin, Germany). A Poisson ratio of 0.5 was assumed for the calculation of the apparent Young’s modulus.

### Real-Time Deformability Cytometry (RT-DC)

RT-DC measurements were performed as described previously(39). Briefly, single cells were re-suspended in PBS containing 0.63% methylcellulose (Sigma Aldrich, St. Louis, USA) at a concentration of 1 × 10^6^ cells ml^−1^. The cell suspension was drawn into a syringe and connected to a microfluidic chip containing two reservoirs connected by a 300-μm- long channel (constriction) with a 30 μm × 30 μm cross-section. The microfluidic chip was made of poly(dimethylsiloxane) (PDMS; Sylgard 184, VWR, Darmstadt, Germany) at which the bottom was sealed with a glass slide (Hecht, Sondheim, Germany). Using a syringe pump cells were driven through the constriction at a constant flow rate of 0.405 μL sec^−1^. Images were acquired at the end of the constriction using a CMOS camera (MC1362, Mikrotron, Unterschleissheim, Germany) which was connected to an inverted microscope (Axiovert, Carl Zeiss AG, Oberkochen, Germany). In real-time cell cross-sectional area (size) and deformation were computed and plotted against each other. It has to be emphasized that deformation and size are not independent parameters for RT-DC. This implies that for two cells of identical mechanical properties, the larger cell will always deform more. A numerical model combining Stokes fluid dynamics with linear elasticity allows to disentangle the relationship of size and deformation and to deduce elastic properties, namely young′s modulus (Mokbel. 2007) Statistical analyses were carried out using 1-dimensional linear mixed model that incorporates fixed effect parameters and random effects to analyze differences between cell subsets and replicate variances, respectively. p values were determined by a likelihood ratio test, comparing the full model with a model lacking the fixed effect term.

### Microfluidic microcirculation mimetic

Microfluidic microcirculation mimetics (MMM) was used including minor modifications as described previously(43). In short, MMM is based on a microfluidic device produced using PDMS and standard photolithographic techniques(29). The microfluidic chip contained an inlet and an outlet connected by a single channel (constriction) with 5 μm smallest in width (maximum width 15 μm). For MMM measurements, single cells were resuspended to a final density of 3 × 10^4^ cells ml^−1^ in PBS containing 0.1% pluronic acid F-127 (Molecular Probes Inc., Eugene, USA). Up to 50 μL of cell suspension was filled into a custom-cut pipette tip (Eppendorf, Hamburg, Germany) and connected to the inlet region of the microfluidic chip. Using a computerized air pressure control system (MFCS-FLEX; Fluigent, Villejuif, France) connected to the inlet and outlet of the device cells were driven through the constriction at a constant pressure of 200 mBar. The microfluidic chip was mounted on the platform of an inverted microscope, which was connected to a camera (The Imaging Source, Bremen, Germany) to record videos at a frame-rate of 120 frames sec^−1^. The first constriction entry time and the total passage time were extracted from the videos using custom-made written codes in MATLAB (The MathWorks, Natick, MA).

### Animal experiments

Animal experiments were strictly performed in compliance with the animal experiment permission no. DD24.1-5131/394/14, approved by the Landesdirektion Sachsen. A total of 1 × 10^6^ mesenpshere MSCs or plastic-adherent MSCs were intravenously injected into the tail vein of NOD scid gamma mice (NOD.Cg-*Prkdc*^*scld*^ *Il2rg*^*tm1Wjl*^/SzJ). After 15 min, organs were resected and used for DNA isolation.

### RNA isolation, complementary DNA synthesis, DNA isolation and real-time quantitative PCR

Total RNA was isolated by TRIzol Reagent (Ambion GmbH, Kaufungen, Germany) and transcribed into complementary DNA using RevertAid First Strand cDNA Synthesis Kit (Thermo Fisher Scientific, Waltham, USA). Real-time quantitative PCR was performed with the primers listed in STable 1 and SYBR Green Master Mix (Thermo Fisher Scientific, Waltham, USA) in a Taqman 7000 Fast cycler (Applied Biosystems, Forster City, USA).

DNA was isolated using DNeasy Blood & Tissue Kit (Qiagen, Hilden, Germany) according to the protocols of manufacturers. Real-time quantitative PCR for Alu sequences was performed as described previously(60). Briefly, 25 μl Taqman PCR Universal Master Mix (Applied Biosystems, Forster City, USA), 900 nM each of the forward and reverse primers, 250 nM TaqMan probe (Table S1), and 200 ng target template were incubated at 50°C for 2 min and at 95°C for 10 min followed by 40 cycles at 95°C for 15 secs and 60°C for 1 min.

All real-time PCR assays were performed in duplicates. The obtained results were corrected to the transcript level of an internal reference gene (glyceraldehyde 3-phosphate dehydrogenase (GAPDH) or hypoxanthine phosphoribosyltransferse 2 (HPRT2)) to yield ΔCT and normalized to control conditions to generate ΔΔCt. The relative change in expression was calculated using the formula 2^-^ΔΔCt.

### Statistical analyses

Despite RT-DC data all statistical analyses were performed using GraphPad Prism Version 5.0 (San Diego, CA, USA).

## Acknowledgements

The authors would like to thank Katharina Rießner, Martina Kalupa, Olaf Penack, and Marta Urbanska for technical support and material provided. Katrin Müller, Kristin Heidel, Felix Reichel, Bärbel Löbel, and Robert Kuhnert for technical support. Microfluidic chips were manufactured with help of the CMCB Microstructure Facility (in part funded by the European Fund for Regional Development-EFRE). Financial support from the DFG SFB655 grant ‘From cells to tissues’ (subproject B2 to C.W. and M.B.), Alexander von Humboldt Stiftung (Alexander von Humboldt professorship to J.G.), Sächsisches Ministerium für Wissenschaft und Kunst (TG70 grant to O.O. and J.G.), an ERC Starting Grant (starting grant “LightTouch” #282060 to J.G.), and DKMS “Mechthild Harf Research Grant” (DKMS-SLS-MHG-2016-02 to A.J.) are gratefully acknowledged.

## Author contributions

S.T. and M.K. designed the project outline and carried out most experiments, interpreted results and co-wrote the manuscript. A.J. provided reagents and supported study design and interpretation. A.T. conducted atomic force microscopy experiments and acquired and analyzed data. M.H. conducted and performed microcirculation mimetic experiments and acquired data. R.W. and M.S. conducted and performed immunomodulatory experiments and acquired and analyzed data. O.O. contributed to real-time deformability cytometry experiments and analyzed data. C.L., B.K. and M.W. were involved in study design. M.B. and J.G designed the project outline, interpreted results and co-wrote the manuscript.

